# Open source tools for temporally controlled rodent behavior suitable for electrophysiology and optogenetic manipulations

**DOI:** 10.1101/243469

**Authors:** Nicola Solari, Katalin Sviatkó, Tamás Laszlovszky, Panna Hegedüs, Balázs Hangya

## Abstract

Understanding how the brain controls behavior requires observing and manipulating neural activity in awake behaving animals. Neuronal firing is timed at millisecond precision. Therefore, to decipher temporal coding, it is necessary to monitor and control animal behavior at the same level of temporal accuracy. However, it is technically challenging to deliver sensory stimuli and reinforcers as well as to read the behavioral responses they elicit with millisecond precision. Presently available commercial systems often excel in specific aspects of behavior control, but they do not provide a customizable environment allowing flexible experimental design while maintaining high standards for temporal control necessary for interpreting neuronal activity. Moreover, delay measurements of stimulus and reinforcement delivery are largely unavailable. We combined microcontroller-based behavior control with a sound delivery system for playing complex acoustic stimuli, fast solenoid valves for precisely timed reinforcement delivery and a custom-built sound attenuated chamber using high-end industrial insulation materials. Together this setup provides a physical environment to train head-fixed animals, enables calibrated sound stimuli and precisely timed fluid and air puff presentation as reinforcers. We provide latency measurements for stimulus and reinforcement delivery and an algorithm to perform such measurements on other behavior control systems. Combined with electrophysiology and optogenetic manipulations, the millisecond timing accuracy will help interpret temporally precise neural signals and behavioral changes. Additionally, since software and hardware provided here can be readily customized to achieve a large variety of paradigms, these solutions enable an unusually flexible design of rodent behavioral experiments.

## Introduction

Mechanistic insight about brain function often comes from accurate recordings of action potentials fired by neurons at millisecond order temporal precision. For instance, to understand how specific information about external stimuli and internal variables is encoded in neural firing patterns, researchers have to measure action potentials from awake, behaving animals (Kayser et al., 2010; Panzeri et al., 2017; Tiesinga et al., 2008; Zuo et al., 2015). Moreover, to have sufficient statistical power to resolve key features of the neural code, animals have to repeat these tasks multiple times in a single recording session (Panzeri et al., 2017). Thus the workflow in behavioral neurophysiology labs typically has two stages: (ii) train animals on a behavioral task of interest and (ii) record their neuronal activity while they perform the task.

The high temporal precision of neuronal firing has strong implications for the scope of suitable behavioral tasks. Intuitively, one can only hope to extract specific information about external variables carried by spike timing (Arabzadeh et al., 2006; Panzeri et al., 2017) if behaviorally relevant events of the task, like cue stimuli (Brunton et al., 2013; Hanks et al., 2015; Jaramillo and Zador, 2011; Ranade and Mainen, 2009; Raposo et al., 2012), go and stop instructions (Lin and Nicolelis, 2008), reward and punishment delivery (Cohen et al., 2012; Hangya et al., 2015; Pi et al., 2013) are under the same precision of temporal control. This intuition is formally captured by Shannon’s framework of information channel capacity, also known as the ‘Shannon limit’, which quantifies the maximal rate of error-free information transfer through a channel with given bandwidth and noise (Csiszár and Körner, 2011).

The first step of analyzing awake recordings is often the calculation of event-aligned spike rasters and different forms of peri-event firing averages (Endres et al., 2008; Shimazaki and Shinomoto, 2010). In these, neural activity is averaged across multiple trials aligned to reoccurring events to provide a statistically robust mean firing activity correlated with the event of interest (Figure 1). This can then be used for instance to interpret the impact of external stimulation on firing rate (Lima et al., 2009) (peri-stimulus time histogram; Figure 1A), to find behavior correlates of specific cell types (Cohen et al., 2012; Hangya et al., 2015; Kvitsiani et al., 2013) (Figure 1B) or to quantify the information about sensory events conveyed by neural activity (Arabzadeh et al., 2006; Diamond et al., 2008; Zuo et al., 2015) (Figure 1C). However, this approach is limited by the available timing information of the aligning event: any unobserved variation of event timing adds to the spiking noise and is mathematically inseparable from biological variation in neural activity (Figure 2). Therefore it is paramount to control and observe the aligning events to the maximal possible accuracy, at least matching the temporal precision of spike timing, necessitating millisecond order precision.

**Figure 1:**
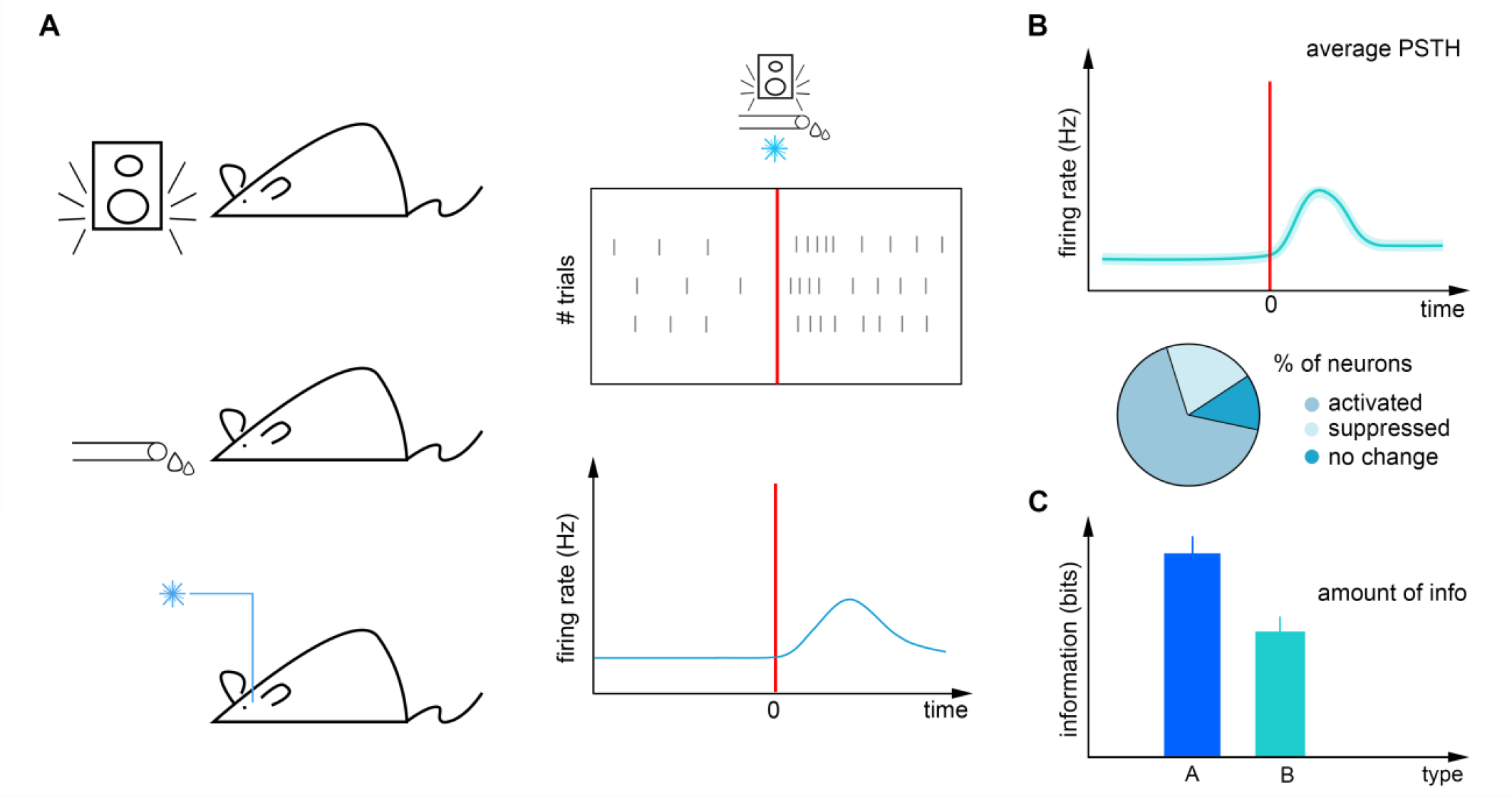
Firing rate analysis. **(A)** Left, mice are exposed to temporally controlled sensory cues, reinforcers and external neural stimulation. Right, neural activity (black ticks) can be aligned to these events in raster plots (top) and peri-event time histograms (bottom). **(B)** Neurons can be categorized according to their tuning to these events. **(C)** The information carried by neural responses about external events can be quantified by information theory tools.

**Figure 2:**
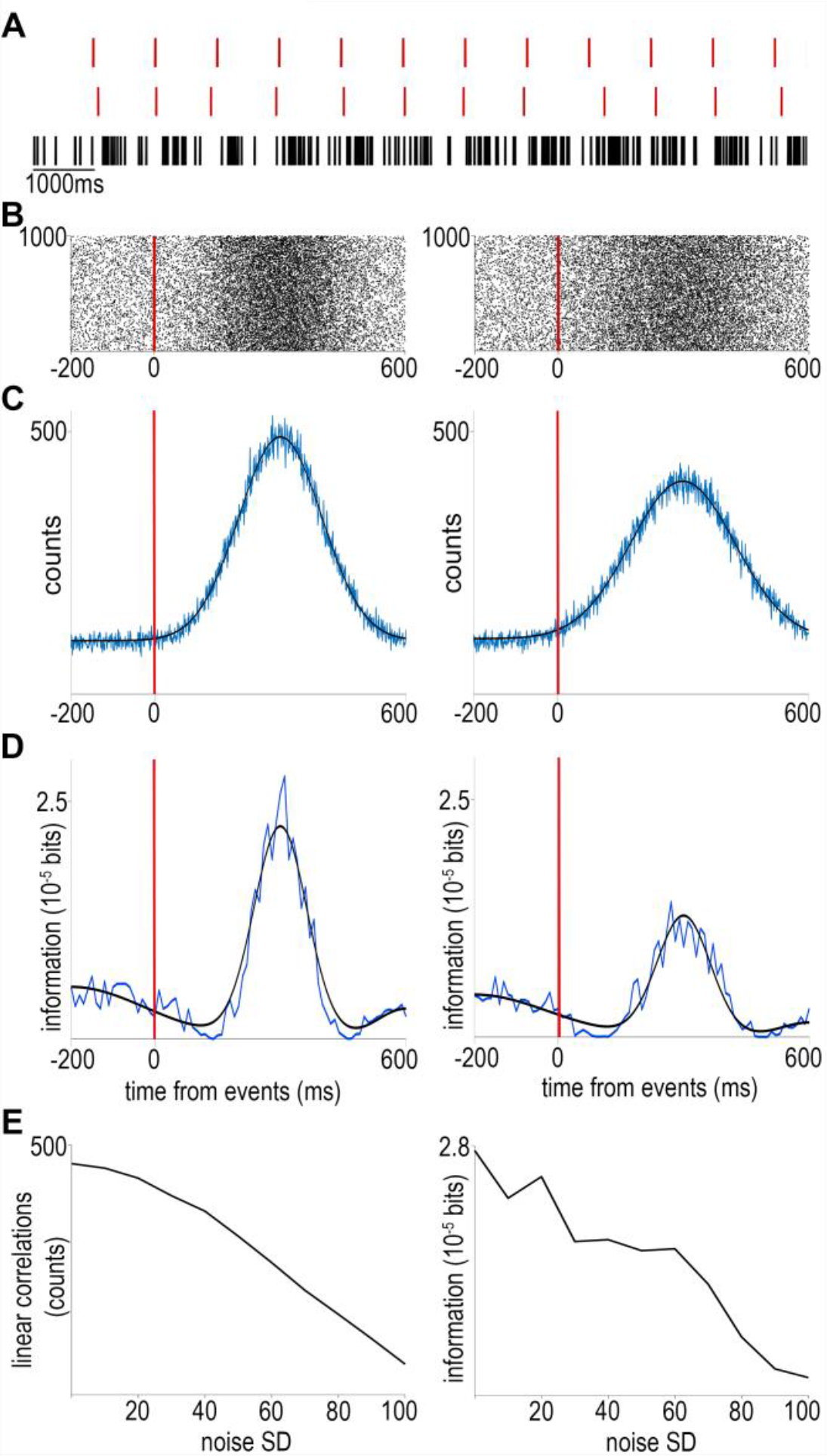
The effects of noise on event timing. **(A)** We simulated an event train without (top) or with Gaussian noise (middle) as well as a spike train consisting of ‘stimulus-evoked’ spikes according to a normal distribution and background Poisson spiking (bottom). **(B)** Raster plots aligned to noiseless (left) or noisy (right) events. **(C)** Peri-event time histograms (equivalent to event-spike cross-correlations) corresponding to the raster plots above. Blue, raw traces; black, Gaussian fits. **(D)** Mutual information between spike times and event times without (left) or with (right) noise. Note the second increase of mutual information around 500 ms corresponding to the information on the event times carried by the lack of event-aligned spikes, demonstrating the power of information theory to detect non-linear correlations. **(E)** Both linear cross-correlation (left) and non-linear mutual information (right) decreases with the amount of added noise in event timing. Code is available at https://github.com/hangyabalazs/Rodent_behavior_setup/, Experiment_simulation.m.

However, standard operating systems and software are not suitable for this task (but see http://rtxi.org/). Indeed, the time from command to task execution varies between times orders of magnitude longer than desired and is influenced by background processes like scheduled system maintenance or security scans. Real-time operating systems can be used to achieve submillisecond behavior control (Brunton et al., 2013; Jaramillo and Zador, 2011; Poddar et al., 2013). However, the emergence of affordable microcontrollers that are easy to program provide a unique opportunity to build modular, flexible and cheap behavior control systems suitable for millisecond order precision (D’Ausilio, 2012; Schultz and van Vugt, 2016).

Many companies offer commercial systems for rodent behavior; however, only few of these are suitable for combining with electrophysiology, typically operating with a limited set of preprogrammed tasks. While some of these provide measurements on noise introduced by parts of the behavioral apparatus, they generally lack delay measurements essential for interpreting neural signals. We developed a behavioral setup capable of temporally precise delivery of sensory cues and reinforcers, demonstrated by measured delay distributions in the millisecond range. Our setup is suitable for combined behavior, electrophysiology and optogenetics experiments. In addition, we describe a custom fabricated sound attenuated behavioral chamber and provide an algorithm for generating calibrated auditory stimuli.

## Methods

### Head-fixed setup

The setup is assembled from a combination of Thorlabs and custom parts. A 3D-printed lick port housing an LED, an infrared photodiode and corresponding infrared photosensor (https://sanworks.io/shop/viewproduct?productID=1010) is mounted on an xyz stage (DT12XYZ/M, Thorlabs). The mouse sits on 3D-printed (https://github.com/hangyabalazs/Rodent_behavior_setup/, stage_rect.skp) rectangular, walled stage mounted on a lab jack (S63081, Fisher). Water delivery for reward and facial air puff are controlled by fast solenoid valves (LHDA0531115H, Lee Company). We use the open source Bpod behavioral control system (Sanworks, https://sanworks.io/shop/viewproduct?productID=1014) for real-time behavioral control and monitor the animals through infrared cameras (FL3-U3-13S2M-CS, Point Grey) using Bonsai open source computer vision software (http://www.open-ephys.org/bonsai/, (Lopes et al., 2014)) (Figure 3A-B).

**Figure 3:**
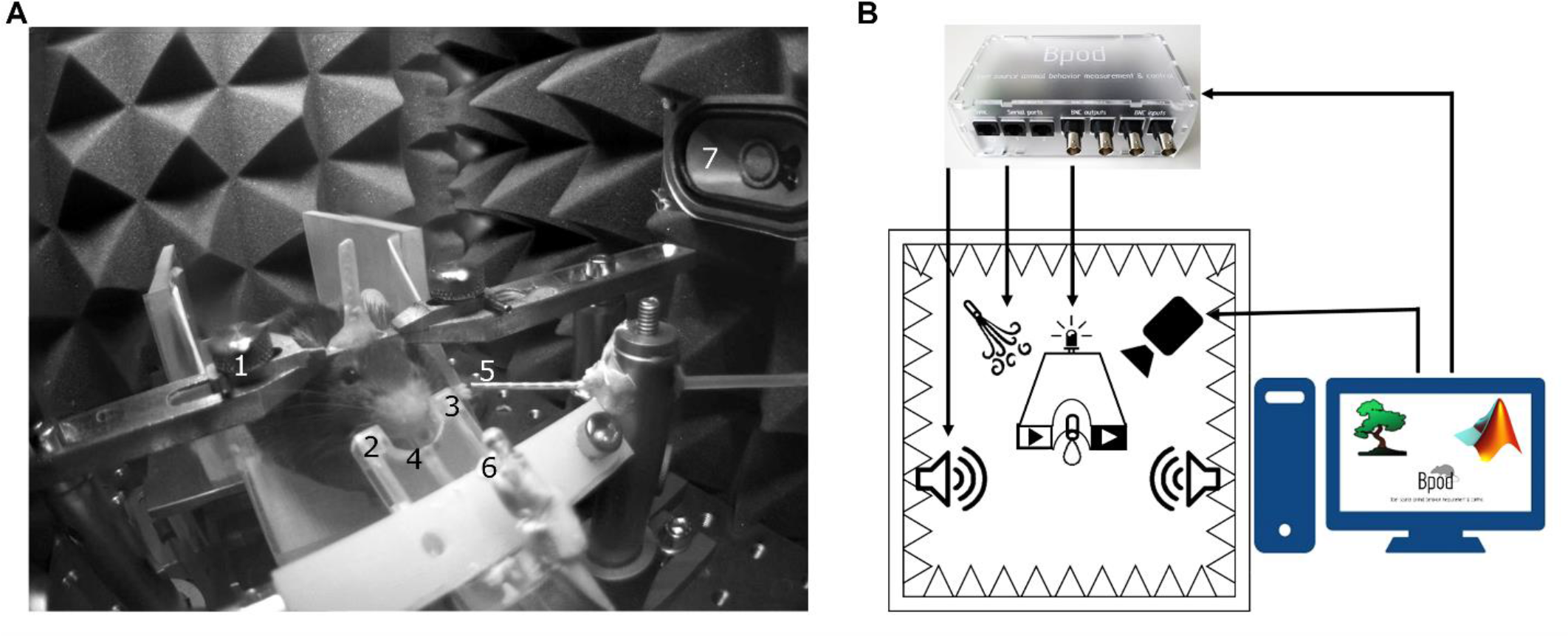
Head-fixed setup. **(A)** The animal is held by an implanted head bar with a pair of metal holders (1), facing a custom-made lick port hosting an IR emitter and an IR receiver (2, 3) for lick detection and a plastic water spout (4). Air-puff is delivered via a cannula placed near the animal’s face (5). Visual and auditory cues are delivered by a central LED (6) and lateral speakers (7). **(B)** Schematic diagram of the behavior setup. Cue and reinforcement delivery are controlled by Bpod. Motion is monitored with a camera using Bonsai open software.

### Sound Calibration

Pure tones were generated in Matlab as sine waves. The tones were uploaded as .wav files to a USB-based microcontroller development system Teensy 3.2 and its audio adaptor board (TEENSY32 and TEENSY3_AUDIO, PJRC) using the Bpod r0.5 behavior control system (Sanworks LLC, www.sanworks.io). The uploaded tracks were played applying Bpod commands controlled by custom-written MATLAB code (https://github.com/hangyabalazs/Rodent_behavior_setup/tree/master/sound_calibration). The Teensy adaptor board’s jack output was connected to an Adafruit audio amplifier Stereo 20w Class D (MAX9744, Adafruit) attached to 8 Ohm Magnetic Speakers (668-1447-ND, Digikey) positioned on the left and right side of the behavioral chamber. A calibrated precision electret condenser microphone (EMM-6, Daytonaudio) was connected to a preamplifier digital converter (AudioBox iOne, PreSonus) and placed in the behavioral enclosure to model the position of the animal’s head in a head-fixed configuration. The sound pressure level (dB SPL) values registered by the microphone were read using the free version of TrueRTA software, a commercial audio analysis software package (TrueRTA, True Audio), and fed back to the custom developed calibration software (Figure 5A-B).

### Measuring the delay of visual cues

LEDs were used for visual stimuli. These were controlled by the Bpod behavioral control system through an open source printed circuit board (‘port interface board’, https://sanworks.io/shop/viewproduct?productID=1008). Stimulus intensity can be directly modulated from Bpod on a 1 to 255 scale. All delays were measured by sending command signals from Bpod on two different outputs: the BNC output port and the RJ45 connector for communication with the port interface board. First, we measured the ‘internal’ delay in addressing these two ports, i.e. the average minimal temporal difference between sending signals on these connectors by Bpod (Figure 6A). The time difference between the two signals was of 0.045 ± 0.001 ms (mean ± SD, n = 180 repeats; see Figure 7A for distribution). This 45 microsecond delay adds only negligible noise to our delay measurements that are on the millisecond order; therefore, we refer to the two signals as ‘concurrent’ hereinafter.

To determine the temporal delay between the command- and the voltage signal sent directly to the LED, a PC oscilloscope (PicoScope 2204A, Pico Technology) was connected to the LED output wire terminal of the port interface board (Figure 6B). Concurrent with the command signal to the board, a TTL pulse was sent to the oscilloscope from the Bpod BNC output terminal. We determined the distribution of temporal differences between the above two signals using oscilloscope measurements (n = 180 repeats).

### Measuring the delay of sound delivery

Sounds were delivered using an audio adaptor board (Audio Adaptor Board for Teensy 3, PJRC) and a microcontroller development system (Teensy USB Development Board 3.2, PJRC) connected to Bpod. To measure the time delay between the command signal and the digital sound signal, the PC oscilloscope was connected to the line out pins of the Audio Adaptor Board (Figure 6C). Whenever a tone was triggered, the signal was detected by the oscilloscope. A BNC cable connected the Bpod module BNC output channel directly to the oscilloscope: the commands for triggering a sound and for sending a TTL pulse were elicited simultaneously and the time difference between the board output and the Bpod TTL was measured as the sound delivery delay (n = 180 repeats).

### Measuring the delay of reinforcement delivery

Air was supplied from a pressurized tank adjusted by an Air Pressure Reducing valve. Water was delivered by gravity from a reservoir placed atop the cage. Reinforcer delivery was controlled by 12V solenoid valves (LHDA0531115H, The Lee Company) connected to the Bpod port interface board. The air and water reservoirs were connected to the valves by Nalgene 180 Clear Plastic PVC Metric Tubing (ID, 2 mm; OD, 4 mm; Thermo Scientific 8001-0204); the same tubing was used between the valves and the lick port. The lick port was equipped by small pieces of polyethylene tubing (ID, 1.14 mm; OD, 1.57 mm; Warner Instruments 64-0755/PE-160) on the output side. The PC oscilloscope was connected to the LED output wire terminal of the port interface board, but the circuit was kept open leaving a small gap (less than 1mm) between the wire from the port interface board and the one connected to the oscilloscope input. The end of the water delivery tubing was placed directly to the gap. Power output was constantly provided to the LED terminal, so upon water outflow the circuit closed and a voltage change could be detected (Figure 6D). To measure the water delivery delay, a TTL pulse was sent to the oscilloscope from the Bpod BNC output terminal in parallel with the logic pulse to the port interface board. The delay was the difference between the signal caused by closing the circuit via the LED terminal and the TTL sent from the Bpod BNC output. The delay of air puff delivery was measured similarly but the circuit through the LED terminal of the port interface board was closed by a small drop of water before measurement and the outflow of air disconnected this circuit. Delay was measured as the time difference between the drop of the signal through the port interface board and the rising edge of the TTL from the Bpod BNC output (n = 60 repeats for each reinforcer).

### Animals and surgery

Electrophysiology and optogenetic stimulation data in this study was obtained from three adult male mice (2 ChAT-IRES-Cre, B6129F1 and 1 PV-IRES-Cre, FVB/AntFx) and behavioral data was presented from an adult male ChAT-IRES-Cre mouse.

For virus injection and microdrive and headbar implantation, mice were anesthetized with an intraperitoneal injection of ketamine-xylazine (0.166 mg/kg and 0.006 mg/kg respectively) after a brief induction with isoflurane. After shaving and disinfecting the scalp (Betadine), the skin was infiltrated with Lidocaine and the eyes were protected with eye ointment (Laboratoires Thea). The mouse was placed in a stereotaxic frame and its skull was leveled along both the lateral and the antero-posterior axes. The skin, connective tissues and periosteum were removed from the scull and a cranial window was drilled above the ventral pallidum / substantia innominate / horizontal diagonal band region of the basal forebrain (VP/SI/HDB, antero-posterior 0.75 mm, lateral 0.6 mm). Cre-dependent Adeno-associated virus (AAV 2/5. EF1a.Dio.hChR2(H134R)-eYFP.WPRE.hGH) was injected into the VP/SI/HDB at 5 mm and 4.7 mm depth from skull surface. Two additional holes were drilled above the parietal cortex for ground and reference. After the virus injection a custom-built microdrive (Hangya et al., 2015; Kvitsiani et al., 2013) was implanted into the VP/SI/HDB using a cannula holder on the stereotactic arm. The microdrive and headbar were fixed with dental cement (LangDental acrylic powder and liquid resin).

### Electrophysiological measurement and optogenetic manipulation

During the surgery, 8 tetrode electrodes (PX000004, Sandvik) were implanted into the VP/SI/HDB along with an optic fiber. Data acquisition was conducted with an Open Ephys board, digitized at 30 kHz. Behavioral data was collected using Bpod and synchronized with the neural data using Open Ephys sync board. For optogenetic stimulation of Channelrhodopsin-expressing neurons in the VP/SI/HDB, we used 1 ms blue laser pulses (Sanctity Laser, SSL - 473 - 0100 - 10TM - D – LED) at 20 Hz frequency with a 2 s on 3 s off duty cycle triggered by PulsePal (1102, Sanworks).

### Sound attenuated enclosure

We used sound absorbing foams (Hanno Sealing and Insulation Systems) designed for machine and commercial vehicle industries. In these foams the sound absorbing element is a combination of an open-cell polyurethane foam and a 25 µm surface skin coated with black synthetic fiber (Hanno Protecto product line, Techfoam). These foams were optimized for airborne sound absorption by converting sound energy to heat as a consequence of friction in the polyurethane cell framework. In our experience, these foams are pliable, easy to cut, handle and mount on vertical surfaces aided by self-adhesive coating. Sound absorbing foams were compared to a 15 mm acoustic insulation board (15mm Acoustic Board, PhoneStar) used for sound insulating floors, walls and ceilings in the construction industry. The acoustic insulation board consists of a fluted cardboard shell and compacted quartz sand filling. The oscillation of loose sand grains converts acoustic energy into kinetic energy. According to specifications, the acoustic insulation board reduces airborne sound by 36 dB and impact sound by 21 dB. However, the boards are relatively heavy (18 kg/m^2^) and there is some sand leakage after cutting to size. We found that while sound absorbing foams and the acoustic insulation board provided comparable levels of sound attenuation, the foam was easier to work with. Although only tested in the 1 kHz to 20 kHz range, based on our measurements and the industrial specifications we extrapolate that similar sound attenuation levels might be achieved in higher frequency ranges relevant for rodent ultrasonic communication. In addition, we used a 70 mm open-cell pyramid foam borrowed from the music studio and stage equipment industry (215894, Muziker) that absorbs higher frequencies and efficiently reduces resonance and echoes (Figure 8). Design file available at https://github.com/hangyabalazs/Rodent_behavior_setup/, sound_attenuated_box.skp

### Bill of Materials

The complete Bill of Materials is available for download at https://github.com/hangyabalazs/Rodent_behavior_setup (headfixed_setup_BOM.docx).

## Results

### Head-fixed setup for combined behavior, electrophysiology and optogenetics

Head-restrained behaviors are becoming increasingly popular due to highly reproducible behavior paradigms with low variability and ease of electrophysiology and imaging experiments (Guo et al., 2014; Hori et al., 2016; Mayrhofer et al., 2013; White et al., 2016).

We provide here a complete modular head-fixed design for combined behavior, electrophysiology and optogenetic experiments in mice (Figure 3). The combination of a photosensor and a photodiode enable temporally precise registration of beam breakings by the animal’s tongue to provide an accurate readout of the animal’s response to sensory cues and reinforcement (Figure 3A and Figure 4A). Alternative to the inexpensive and widely used infrared sensors (Jaramillo et al., 2014; Lin and Nicolelis, 2008; Shin et al., 2018), solutions relying on the closure of an electric circuit by the animal’s tongue are also available (Marbach and Zador, 2016; Petykó et al., 2015; Slotnick, 2009; Wilson et al., 2017). We opted for the photosensor/photodiode solution in order to avoid the introduction of electric artifacts to the electrophysiology recordings and because the same port structure could be employed for head-fixed and freely behaving paradigms. For head-fixation, a custom pair of metal head bar holders are used. It is important that the mouse is positioned comfortably in the head-restraint and can properly reach the lick port within its natural range of tongue movements (Figure 3A). Indeed, in our experience, the most common mode of failure in training is the inadequate positioning of the licking spout. Since mice differ in size and position of their head restraint implants, two opportunities of adjustment are built in the head-fixed setup. First, the mouse is positioned on a rectangular, walled stage mounted on a lab jack that allows for adjustments in height. Second, the lick port is attached to an xyz stage allowing fine spatial positioning of the licking spout with respect to the animal’s snout – a crucial step for achieving stable head-fixed performance. To achieve temporally precise behavioral feedback, water delivery for reward and facial air puff are controlled by fast solenoid valves. We use the open source Bpod behavioral control system for real-time behavioral control and monitor the animals through infrared cameras using open source computer vision that allows the tracking of eye blinks, licking, whole body movements or pupil diameter (Figure 3B). This setup is suitable for combining head-fixed mouse behaviors with single neuron recording and optogenetic stimulation (Figure 4B-D).

**Figure 4:**
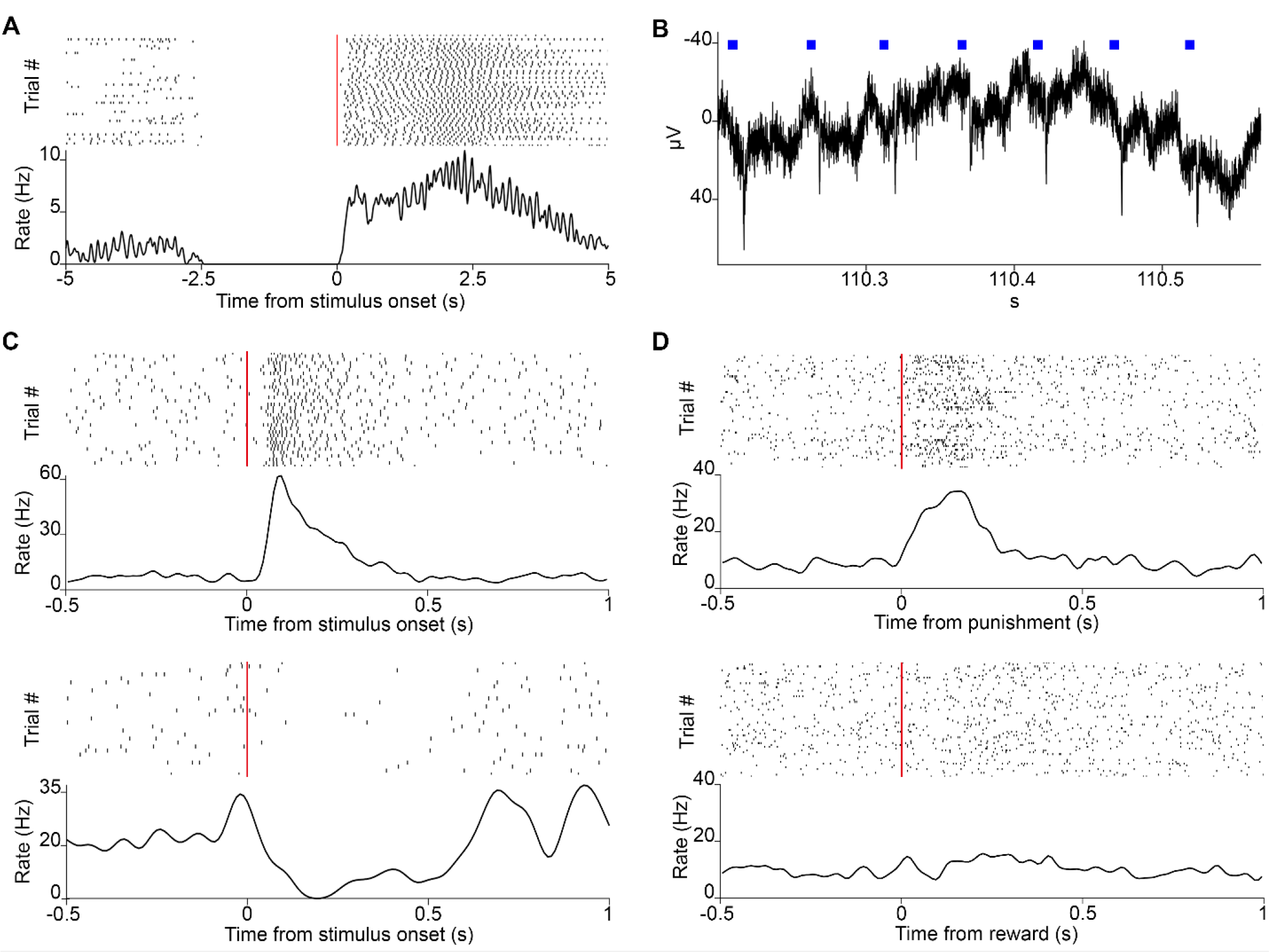
Neuronal recordings. **(A)** Raster plot of lick responses (black ticks) to reward-predicting cues (red line) of a mouse trained on an auditory Pavlovian task. **(B)** Local filed potential deflections in response to photostimulation (blue squares) in a mouse expressing the light-sensitive channelrhodopsin in parvalbumin-expressing neurons of the HDB. **(C)** Raster plot of action potentials (black ticks) of two neurons recorded in VP/SI responding to the reward-predicting cues (red line) in Pavlovian conditioning. **(D)** Raster plot of a neuron from VP/SI selectively responding to punishment but not reward in the same task.

### Delivering calibrated tones

In behavioral experiments it is often important to parametrically control stimulus intensity, for instance to manipulate task difficulty. We developed a sound calibration protocol to precisely control the sound pressure level (dB SPL) of auditory stimuli that allows both playing tones of controlled intensity and sound equalization.

A precision microphone was connected to a preamplifier digital converter and placed between the head bar holders to model the position of the animal’s head in the head-fixed setup (Figure 5A). Pure sinusoidal tones from 1 to 21 kHz frequency (with 0.25, 0.5 or 1 kHz resolution) with a predefined arbitrary amplitude were loaded onto the Teensy board via Bpod and played in sequence. Actual dB SPL levels of the played tones were measured with the calibrated microphone using the free version of the spectrum analyzer software TrueRTA and fed back to the calibration protocol software (Figure 5B). The experimenter then selected the target dB SPL and the amplitudes of the sine waves were updated according to Equation 1:

**Figure 5:**
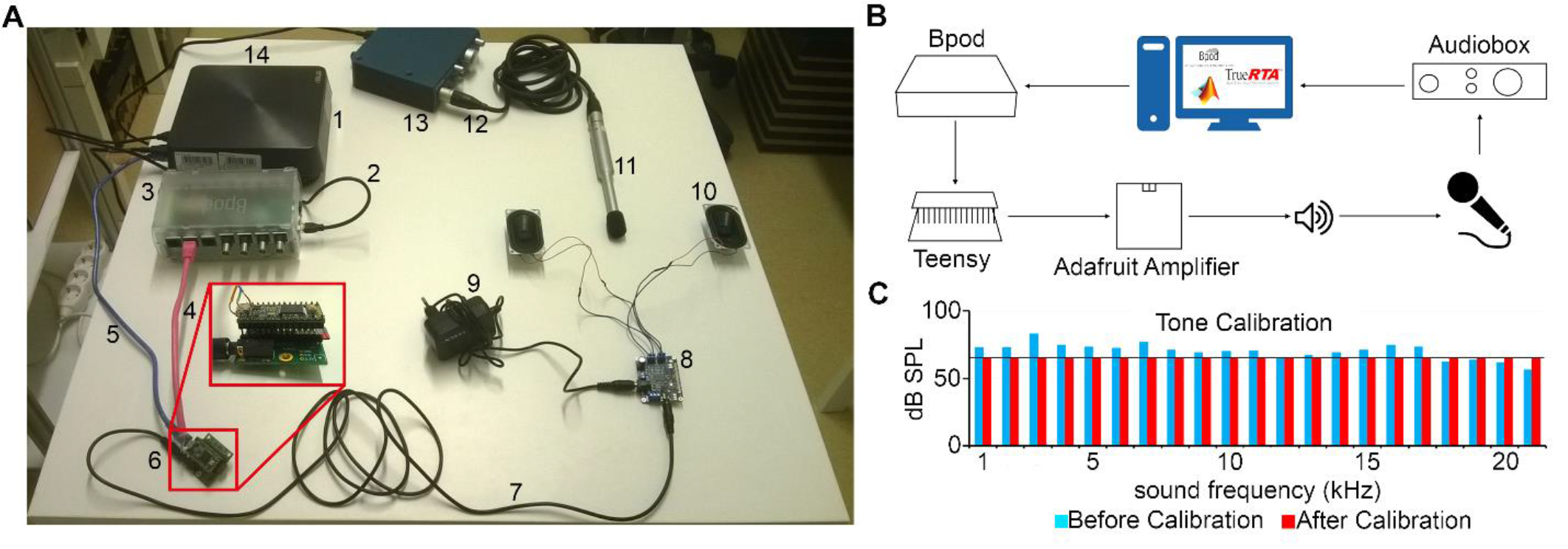
Sound calibration and delivery. **(A)** Components: computer (1), miniUSB-USB A cable (2), Bpod (3), RJ45 cable (4), miniUSB-USB A cable (5), Audio Adaptor Board for Teensy + Teensy USB Development Board + SD card (6), 3.5mm stereo jack to jack cable (7), Adafruit Audio Amplifier (8), 12V power supply (9), Digikey 8 Ohm Magnetic Speakers (10), EMM-6 Electret Measurement Microphone (11), Male-Female three-pin XLR cable (12), AudioBox iOne (13), USB B -USB A cable (14). **(B)** Schematic of the setup. A sine wave is generated in Matlab and sent to Bpod, which loads it to the Teensy apparatus as a .wav file. When played by the speakers, the sound is detected by the microphone, delivered to the computer and the dB SPL is read by the TrueRTA software. **(C)** The dB SPL levels at each frequency before (blue) and after (red) the calibration process. Solid black line indicates the calibration target volume (60 dB SPL).

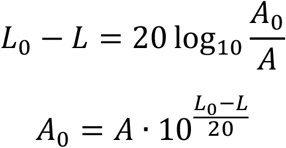

where *L* is observed and *L_0_* is the target dB SPL, *A* is the actual sine wave amplitude and *A_0_* is the target amplitude corresponding to *L_0_*.

Based on this calibrated set of sound intensities, any target intensity can later be software generated using Equation 1 (Figure 5C). It should be noted that this protocol does not handle sounds above 21 kHz, since common commercially available speakers cannot efficiently play sounds above the human hearing range. However, provided specific equipment for ultrasounds is available, the code can easily be adjusted to the experimenter’s needs.

Besides pure tones, the calibration protocol also produced calibrated white noise. The procedure was similar to the pure tone calibration described above. White noise (*WN*) was generated with a pre-assigned amplitude (*A* = 0.2) using Equation 2:

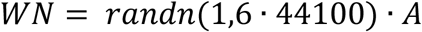

*A_0_* was calculated for five separate frequencies (1, 2, 5, 10 and 20 kHz) using Equation 1 and their average was used to assign the new amplitude for *WN*. We would like to note that generating equalized white noise by mixing calibrated frequency components would also be possible.

### Temporally precise delivery of visual and auditory stimuli

In order to correlate action potential firing of neurons with external stimuli, it is important to accurately control the timing of stimulus presentation. We demonstrate a method to deliver simple visual and auditory stimuli to mice with millisecond order temporal precision (see Materials and Methods; Figure 6). We measured the exact latency and jitter of stimulus presentation. It is important to note that while the jitter directly increases the measurement noise for action potential timing (Figure 2), measured mean latencies can be compensated for post hoc.

**Figure 6:**
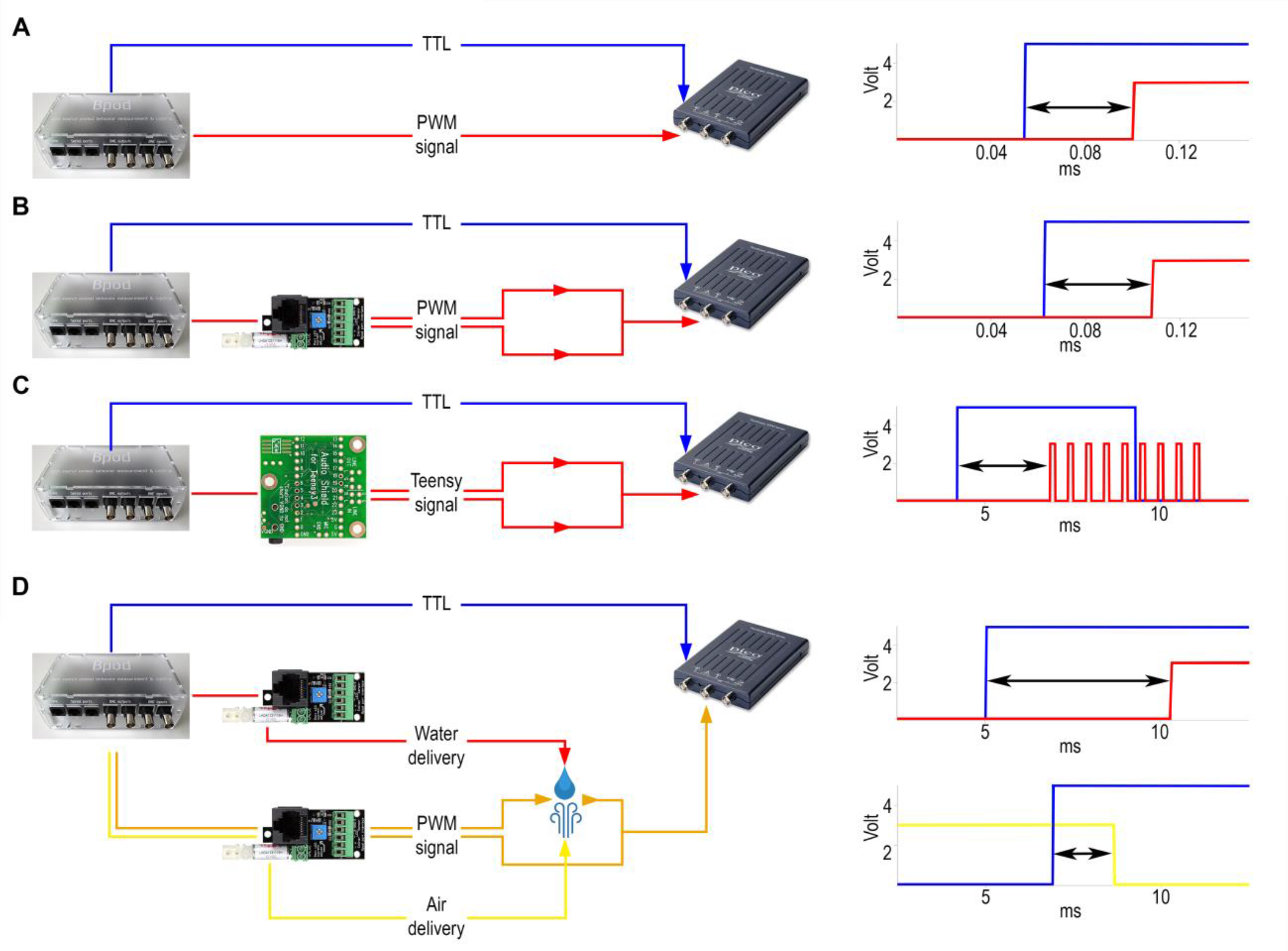
Delay measurements. **(A)** Internal delay. Left, signals were sent from the BNC output port (blue) and the RJ45 output connector for communication with the port interface board (red) directly to the oscilloscope. Right, example signals detected by the oscilloscope. Arrow, measured delay **(B)** Delay of visual cue. Left, signals were sent from Bpod to the oscilloscope both directly (blue) and via the port interface board (red). Right, example of the signals detected by the oscilloscope. **(C)** Delay of sound delivery. Left, signals were sent directly (blue) or via the Teensy board (red). The oscilloscope receives the latter signal from the line out pins of the Teensy slave board. Right, example of the signals detected by the oscilloscope. **(D)** Delay of reinforcement delivery. Left, signals were sent from the BNC output port (blue) directly to the oscilloscope and to two port interface boards. One port was receiving commands to open and close the water valve (red), while the other was receiving similar input for controlling the air valve (yellow) along with a constant PWM signal (orange). The latter was sent to the oscilloscope throughout a circuit that water or air could close or break, respectively, changing the oscilloscope voltage input. Top right, example of the signals detected by the oscilloscope for water delay measurement. Bottom right, example of the signals detected by the oscilloscope for air delay measurement.

To assess temporal precision of light delivery, we measured the time delay from the commanding TTL signal to the signal coming from the Bpod port interface board LED pins (see Materials and Methods; Figure 6B). Average light latency was 0.047 ± 0.003 ms (mean ± SD) (Figure 7B).

**Figure 7:**
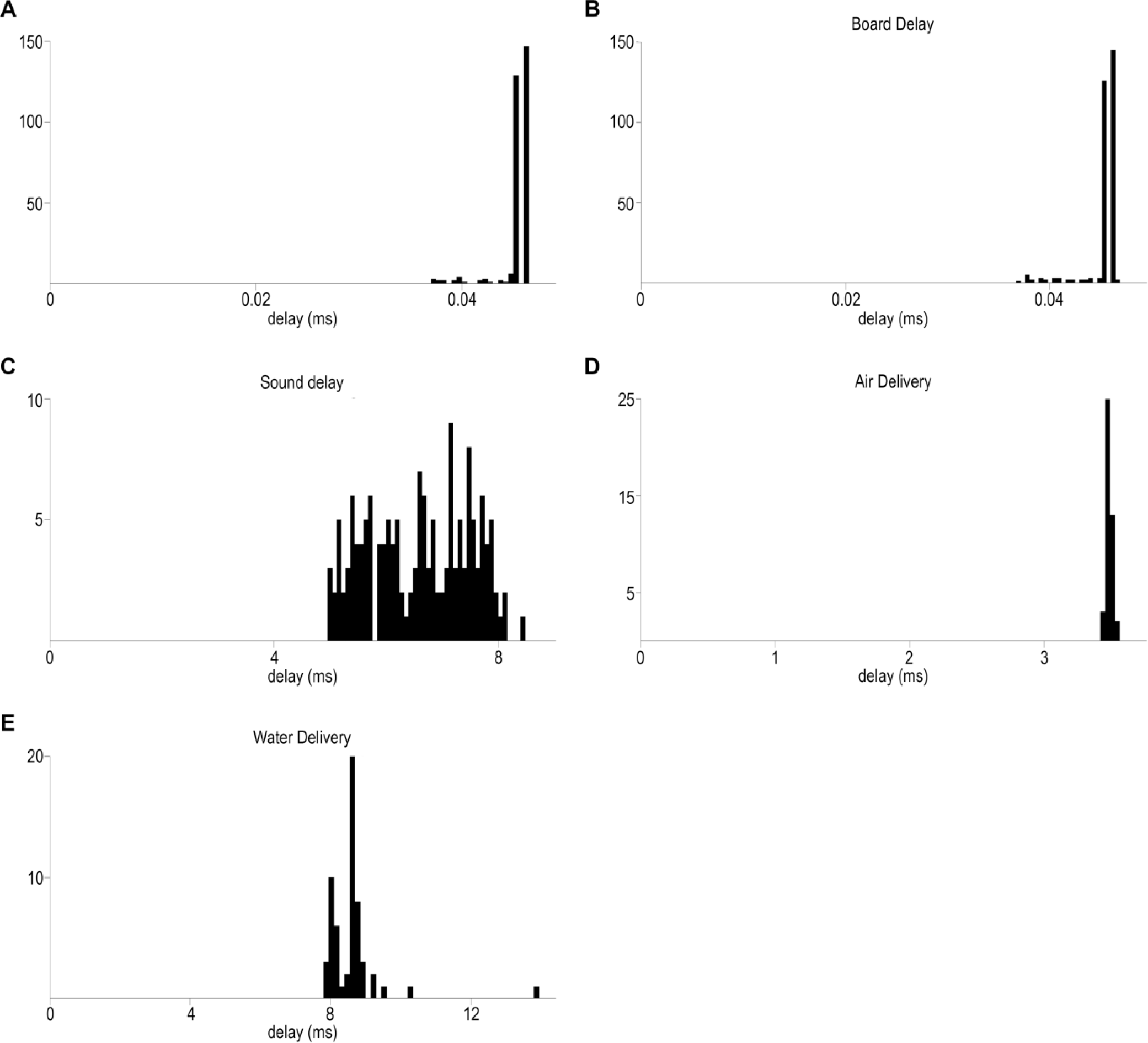
Temporally precise delivery of stimuli and feedback. **(A)** Distribution of minimal elapsed time between sending signals to the BNC and RJ45 output of Bpod (mean ± SD, 0.045 ± 0.001 ms). **(B)** Board delay: distribution of delays between the signals from the BNC output port and the LED output wire terminal of the port interface board (mean ± SD, 0.047 ± 0.003 ms) **(C)** Delay distribution of sound delivery, between the signals from the BNC output port and the Teensy board (mean ± SD, 6.59ms ± 0.9 ms). **(D)** Delay distribution of air puff delivery (mean ± SD, 3.48 ± 0.02 ms). **(E)** Delay distribution of water delivery (mean ± SD, 8.61 ± 0.81 ms).

Temporally precise delivery of complex audio signals with short latency is a more complicated problem, since it requires a sound card that can store digital waveforms and send analog signals to speakers in order to play the sounds. Therefore, we used a microcontroller development system with an audio adaptor board for playing sounds (see Materials and Methods; Figure 6A-B). Pure tones were constructed as sinewaves of characteristic frequencies, with specific sampling rate, time duration and amplitude and uploaded to a memory card. Uploading and playing the sounds was achieved through the behavioral control system. We measured the temporal delay from the TTL command onset to the onset of the analog signal sent from the audio adaptor board to the speakers (Figure 6C). Sound onset latency was 6.59 ± 0.9 ms (mean ± SD) (Figure 7C).

### Temporally precise delivery of reinforcers

In associative learning paradigms, animals are rewarded and punished with appetitive and aversive stimuli, respectively. The temporally precise delivery of these stimuli, hereafter referred to as reinforcers, is paramount if we wish to understand how action potential firing of different neurons is related to these events. We equipped our setup with a system of solenoid valves and silicon tubing in order to deliver water reward and air puff punishment. We optimized these delivery systems to keep the distances short and avoid more bending than necessary, in order to minimize delays in water and air delivery. We note however, that the exact latencies may depend on the tubing configuration, therefore we suggest performing delay measurements for every unique configurations (see details of tubing specifications in Materials and Methods). The opening and closing of valves that allow air and water flow was controlled by TTL pulses triggered by the behavioral control system. To estimate the temporal precision of water and air delivery, we measured the time lags between the onset of the commanding TTL pulses and water or air leaving the tube system at the position of the animal’s face.

We determined the time of water delivery by allowing the water droplet exiting the end of the delivery tubing to close an electric circuit by touching two wires positioned close to each other. Conversely, we measured air puff delivery by allowing the outflowing air wave to break a closed circuit by removing a drop of water from between two closely positioned wires. The rise (water) or fall (air) of the circuit signal was compared to a TTL pulse sent from the behavior control unit concurrent with the valve command signals (Figure 6D). Latency of air delivery was 3.48 ± 0.02 ms (mean ± SD). Delay of water delivery was 8.61 ± 0.81 ms (mean ± SD) (Figure 7D-E).

### Sound attenuated enclosure

For efficient experimentation, rodent behavioral training is typically parallelized such that multiple animals are trained at the same time. This necessitates ‘compartmentalization’, that is, the sensory isolation of the individual experiments. To this end we designed a sound attenuated chamber (Figure 8A-B). The enclosure was built of medium-density fiberboard (MDF) since dense wood fiber is known to effectively reduce both airborne and impact sound. Sound attenuation was achieved by a combination of the dense engineered MDF wood, sound absorbing foams or an acoustic insulation board and pyramid acoustic foams. In order to filter electrical fields for carrying out low noise electrophysiological measurements, the interior of the box was covered with dense stainless steel mesh fabric (Figure 8C).

**Figure 8:**
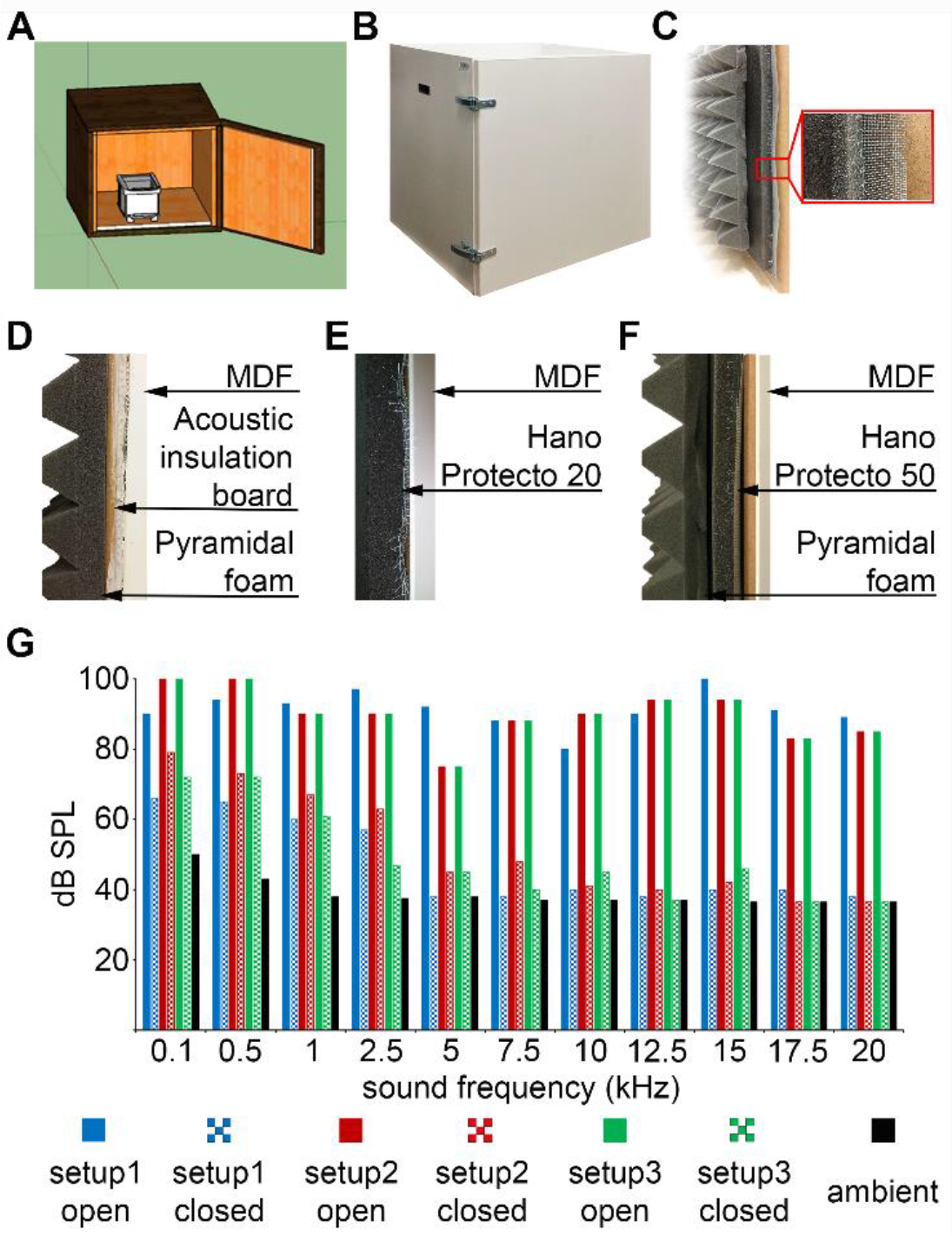
Custom-made sound-attenuated enclosures. **(A)** A 50-by-50-by-50 cm sound attenuated box designed in the freely available 3D modeling software SketchUp. **(B)** Picture of the sound attenuated chamber. **(C)** Cross-section of the box: from left to right, pyramidal foam, sound absorbing foam, stainless steel mesh and medium-density fiberboard (MDF). **(D)** Configuration #1: sound attenuation by acoustic insulation board with quartz sand filling and pyramidal foam. (E) Configuration #2: Hanno Protecto 20 foam. **(F)** Configuration #3: Hanno Protecto 50 foam combined with pyramidal foam. **(G)** Sound attenuation measurements: pure tones of different pitch were played from speakers outside the enclosure and the dB SPL was measured by a microphone placed inside with the door open or closed.

We tested three different configurations for sound attenuation (Figure 8D-F) and compared the obtained attenuation levels at different tone frequencies (Figure 8G). Sound pressure level was measures by a calibrated microphone placed inside the behavioral chamber while tones were played by speakers positioned outside the apparatus. We played 80-100 dB pure tones at different frequencies ranging from 100 Hz to 20 kHz outside the enclosure and measured the attenuated sound inside the chamber. From 2 kHz, which corresponds to the lower limit of rodent hearing range (Koay et al., 2002), all three setups reduced the detected volume to 35-45 dB, which did not differ from ambient noise levels (Figure 8G). Thus both the sound insulation board and the sound absorbing foams achieved sufficient sound attenuation (Koay et al., 2002) while the absorbing foams had superior physical qualities (lower weight and no sand leakage).

Behavioral apparatuses for multiple behavioral paradigms can fit inside the sound attenuated enclosure, including both freely moving (e.g. two-alternative forced choice, 5-choice serial reaction time task (Busse et al., 2011; Kim et al., 2016; McGaughy et al., 2002; Yang and Zador, 2012; Yoshida and Katz, 2011) and head-fixed (e.g. operant auditory go/no-go detection and discrimination tasks, Pavlovian conditioning (Cohen et al., 2012; Eshel et al., 2015; Hangya et al., 2015; Pinto et al., 2013; Sanders and Kepecs, 2012)) behaviors.

## Discussion

We presented a modular behavioral training environment to perform well-controlled, high-throughput behavioral assays in rodents. We provided a way of delivering audio-visual stimuli of calibrated intensity and measured the precise temporal latencies and jitters of stimulus and reinforcement delivery in this behavioral training solution. We believe this affordable modular system will allow streamlining behavioral experiments combined with electrophysiology and optogenetic manipulations.

Likely due to their flexibility, sufficient precision, low price and the relatively user-friendly programming options available, there has been a recent increase in microcontroller-based applications in neuroscience (D’Ausilio, 2012; Schultz and van Vugt, 2016). An impressively simple apparatus realizing a classical Skinner box is described by (Pineño, 2014), interfacing an Arduino Uno and an IPod Touch (coined the ArduiPod Box) capitalizing on the tablet touch sensor. However, being a highly specialized solution, the system does not allow the implementation of complex paradigms and using the iOS makes protocol development and interfacing with additional components slightly complicated. A more sophisticated setup based on Raspberry Pi and Python was demonstrated for operant licking behavior and sucrose preference, potentially capable of integrating multiple components (Longley et al., 2017). Rizzi et al. described an Arduino-driven operant box for optogenetic self-stimulation (Rizzi et al., 2016); however, potential extensions of this application are limited by lack of online feedback and controlling tools. This was overcome in another Arduino-based operant chamber capable of delivering water reinforcer (Devarakonda et al., 2016) that provides a GUI for live data monitoring. Versatile options for head-fixed behavioral tasks combined with two-photon calcium imaging have been developed by Micaleff et al. for go/no-go tasks (Micallef et al., 2017) and by Brugess and colleagues for two-alternative choice paradigms (Burgess et al., 2017). Here we provide a complex system that goes one step further in flexibility and level of control compared to these more specialized solutions. While our behavior setup is suitable for the implementation of most head-fixed and freely behaving rodent paradigms in combination with multiple single neuron recording and optogenetic stimulation with live performance feedback through the Bpod GUI, we have rather put our main emphasis on precision control. First, we developed a sound delivery system for calibrated pure tones and white noise and any combinations of these stimuli, in which sound pressure levels can be precisely adjusted. This required the development of an inexpensive version of a sound attenuated enclosure for better isolation from external sources of disturbance. Second, we provided delay measurements for cue stimuli and reinforcement delivery and made significant efforts to minimize these latencies and jitters, achieving submillisecond precision in reinforcement timing. Indeed, very few such measurements has been made available to date (Chen and Li, 2017; D’Ausilio, 2012; Schultz and van Vugt, 2016), and those are mostly restricted to microcontroller board delays or simple cue stimuli.

Of note, a similar approach is starting to be adopted for small non-human primates. The Operant Box for Auditory Tasks (OBAT) (Ribeiro et al., 2017) is placed in a sound-attenuated chamber and capable of delivering auditory cues that marmosets learn to discriminate. Audio traces are stored and played from an SD card with latencies of 4.607 +- 0.326 ms. We measured average latencies of 6.59 ± 0.9 ms with the Teensy system that is capable of delivering mixtures of pure tones and white noise, ideal for auditory detection tasks, as well as other complex stimuli. However if such complex stimuli are not required, cheap mini speakers and piezo buzzers (e.g. Adafruit or Mouser Electronics) are also capable of delivering pure tones of fixed frequencies typically between 400 Hz and 22 kHz and amplitudes controlled by the current passed. With minor modifications these devices can be incorporated in our setup controlled by Bpod, potentially reducing latencies of auditory stimuli to submillisecond levels.

### Temporal control

Precise temporal control of the behaviorally relevant events of the task is essential for reliable estimation of spike timing with respect to those events. Major goals of a behavior control system that provides an ideal basis for electrophysiology were spelled out around the relatively recent birth of the rodent cognition field: “(1) interact rapidly with the experimental subjects […] the system should respond as fast as possible, and do so reliably; (2) provide high-time-resolution measurements of the events that have occurred, so as to get reliable time traces of behavior when combining it with electrophysiology; (3) be flexible yet easy to program and modify” (http://brodywiki.princeton.edu/bcontrol/). These principles lead to the design of BControl, a behavior control system relying on the precision of the Real Time Linux State Machine. This system fulfilled the first two goals really well, and the third one as well at then state-of-the-art. What has changed is the increasing availability of cheap and easy-to-program microcontrollers, which now allow almost the same level of precision but with a considerably less bulky system. Instead of the combination of a Windows machine, a complicated Real Time Linux machine, an expensive data acquisition card and correspondingly complex governing code, a single microcontroller with significantly simpler software interface can fulfill the goals delineated above (Sanders and Kepecs, 2012, 2014).

While we provide a way of delivering primary reinforcement with submillisecond jitters, the precision of reinforcement delivery could be further increased by direct stimulation of reward- or punishment-related areas (Kim et al., 2012; Kravitz et al., 2012) or using secondary reinforcement schemes (Slawecki et al., 1999). However, since the underlying neural mechanisms for processing direct brain stimulation or secondary reinforcement are likely different (Beck et al., 2010), these alternatives will not fully substitute precise delivery of primary reinforcers.

### Spatial control

Head-fixed experimental paradigms has been criticized because the animals are artificially restricted in their natural movements. While true, this also has an often overlooked flip side. Constraining the space of possible behavioral patterns of the animal means better behavioral control at the same time, as it decreases the uncontrolled behavioral variability and hence the uncontrolled variability (‘noise’) of the electrophysiological recordings. In addition, it allows optical imaging of neural activity under fixed microscopes, which has been a strong motivation for head-fixed designs (Chen et al., 2013). Thus these experiments are easily reproducible and offer a good opportunity to investigate neuronal mechanisms in non-anesthetized, behaving animals in a large number of repeated trials within one session (Guo et al., 2014; Micallef et al., 2017). Accordingly, head-fixed experimental procedures are becoming increasingly popular in studying neural mechanisms of sensation, cognition and other behaviors in rodents (Abraham et al., 2012; Dolzani et al., 2014; O’Connor et al., 2010; Peters et al., 2014; Saleem et al., 2017; Yu et al., 2016).

Nevertheless, many of the behavioral paradigms require free motion of the animals. For instance, although there are an increasing number of very useful virtual reality systems for rodents (Harvey et al., 2009; Keller et al., 2012), many of the spatial learning paradigms are still performed in freely behaving animals. In addition, an important line of research is addressing the more naturalistic animal behaviors including, but not restricted to motor learning and execution, in which case minimizing the artificial constraints is inherent to the experiment (Kawai et al., 2015; Lopes et al., 2017). How do we achieve the best possible spatial control in these experiments? We argue that tracking the spatial position of the animal or its specific parts (limbs, joints, head, tail, etc.) to the same level of precision as compared with other aspects of their behavior is crucial. Not only does it allow precise registering of action potential firing with respect to spatial features, it also opens the possibility of closed-loop stimulation based on position. Note that the open source Bonsai software operating on the principle of parallel asynchronous streams is well suited for these tasks (Lopes et al., 2014).

### Concluding remark

We demonstrated a modular, flexible, affordable, largely open source solution for rodent behavior with sufficient temporal precision for electrophysiology and optogenetic experiments. Although we mainly focus on measuring the temporal delays partly originating from a piece of commercial hardware (Bpod, Sanworks LLC), our methods and descriptions are of general use, for the following reasons.

First, in systems neuroscience, correlation of neuronal activity with behaviorally relevant events is of prime interest. Indeed, the construction of raster plots and peri-event time histograms is considered the ‘gold standard’ analysis by many researchers (Figure 1). However, accuracy of alignment between behavioral and neural data is constrained by the precision to which behaviorally relevant events are registered (Figure 2). Therefore, measuring temporal precision of behavior control systems is of broad interest.

Second, only a small fraction of the measured delays originated from Bpod Arduino boards compared to delays in the tubing and valve system for reinforcers and those introduced by the Teensy system and potentially other components for sound delivery (Figure 7). We made significant efforts to minimize these latencies given the constrains set by complex behavioral tasks. Of note, we have provided a way to characterize these delays (Figure 6) that can be implemented in any such systems. Since delay measurements of this type are sparse, we expect that these data will be of interest.

Third, currently only few systems offer combined behavioral training, electrophysiology and optogenetics (Figures 3-4) with high level of flexibility of experimental design. Therefore, we believe that the description of our setup is useful for groups interested in performing such experiments.

Fourth, auditory physiology labs use expensive high-end equipment for sound calibration, typically unavailable in standard electrophysiology labs. We have provided a detailed description of an inexpensive sound attenuating box (Figure 8) and a configuration for sound calibration and equalization (Figure 5), including Matlab code that can be adapted to the experimenter’s needs.

Some labs moved in the direction of automating behavioral training (Erlich et al., 2011) or even electrophysiological recordings from behaving animals (Dhawale et al., 2015). As these systems have clear benefits, we anticipate that they will be tremendously useful in the future. Nevertheless, they require considerably more resources than the simple system presented here. Also, we see the advantage of our system in its near-complete flexibility, which may be more useful during the planning, developing and troubleshooting phases of the experiments. In addition, our system also carries the possibility of multiplying into large scale, automated systems.

We anticipate a continuing growth of the technical armaments of behavioral neurophysiology suited for ever-increasing temporal and spatial control. Sharing new tools within the scientific community is important in order to facilitate this technical development. Design files, Matlab code, bill of materials and other documents pertaining to the present report can be downloaded from https://github.com/hangyabalazs/Rodent_behavior_setup/.

## Ethics Statement

All experiments were approved by the Committee for the Scientific Ethics of Animal Research of the National Food Chain Safety Office (PE/EA/675-4/2016) and were performed according to the guidelines of the institutional ethical code and the Hungarian Act of Animal Care and Experimentation (1998; XXVIII, section 243/1998, renewed in 40/2013) in accordance with the European Directive 86/609/CEE and modified according to the Directives 2010/63/EU.

## Author Contributions

BH developed the idea and conceptualized the manuscript. KS and PH designed the behavioral environment. PH performed combined experiments of behavior, electrophysiology and optogenetics. NS and TL performed the delay measurements. NS developed the sound delivery system and performed all sound measurements. NS and BH performed the simulations. NS and KS generated the figures. BH and NS wrote the manuscript with input from all authors.

## Acknowledgement

We thank Ranade P. Sachin and Rob Eifert for parts of the head-fixing apparatus, Péter Barthó and Sándor Borbély for their input on the sound attenuated enclosure, Joshua I. Sanders and András Széll for help with the Bpod system and the delay measurements, Adam Kepecs for concepts and discussions. This work was supported by the ‘Lendület’ Program of the Hungarian Academy of Sciences (LP2015-2/2015) and the European Research Council Starting Grant no. 715043. BH is a member of the FENS-Kavli Network of Excellence.

